# A nucleotide-sensing endonuclease from the Gabija bacterial defense system

**DOI:** 10.1101/2020.12.18.423425

**Authors:** Rui Cheng, Fengtao Huang, Hui Wu, Xuelin Lu, Yan Yan, Bingbing Yu, Xionglue Wang, Bin Zhu

## Abstract

The arms race between bacteria and phages has led to the development of exquisite bacterial defense systems including a number of uncharacterized systems distinct from the well-known Restriction-Modification and CRISPR/Cas systems. Here, we report functional analyses of the GajA protein from the newly predicted Gabija system. The GajA protein is revealed as an endonuclease unique in that: 1. It may function as a restriction enzyme or a site-specific nicking enzyme, depending on the arrangement of the recognition sequences; 2. Its activity is strictly regulated by nucleotides concentration. NTP and dNTP at physiological concentrations can fully inhibited the robust DNA cleavage activity of GajA. Interestingly, the nucleotide inhibition is mediated by an ATPase-like domain, which usually hydrolyzes ATP to stimulate the DNA cleavage when associated with other nucleases. These features suggested the mechanism of the Gabija defense in which an endonuclease activity was suppressed at normal condition, while activated by the depletion of NTP and dNTP upon the replication and transcription by invaded phages. This work highlights a concise strategy to utilize a single protein for phage resistance via nucleotide regulatory.

## INTRODUCTION

To resist frequent and diverse attacks by bacteriophages, bacteria have developed multiple, exquisite defense strategies that can collectively be referred to as the bacterial “immune system” (1-3). Anti-phage defense strategies include the adaptive immune system CRISPR/Cas, which provides acquired immunity by memorizing past phage invasion (4); innate immune systems restriction-modification (R-M) systems that target specific sequences in the viral DNA (5); abortive infection systems (Abi) systems that cause cell death or metabolic disturbance upon phage infection (6); and additional systems with mechanisms that are not yet clear. In recent years, CRISPR/Cas9 gene editing technology derived from CRISPR/Cas system developed so swiftly and has been the most widely used gene editing method. Similarly, restriction endonucleases derived from R-M systems had led to revolutions of recombinant DNA technology and served as key enzymatic reagents for modern molecular biology. The most widely used restriction enzymes in laboratory are Type II restriction endonucleases, which were further classified into 11 subtypes: A, B, C, E, F, G, H, M, P, S and T based on their property and behavior (7). For example, the enzymes of ‘IIP’ subtype recognize palindromic (symmetric) DNA sequences and the ‘IIS’ subtype enzymes are characterized by shifted cleavage. Pingoud et al. have discussed the mechanisms of sequence recognition and catalysis of Type II restriction endonucleases systematically (8).

Recent booming of metagenomic analysis has suggested that a large number of uncharacterized defense systems exist in bacteria (9). As predicted, an increasing number of defense system were validated successively (10-12). Doron et al. predicted and experimentally verified ten potential anti-phage defense systems, although their molecular mechanisms are not yet understood (10). Recent progress on the underlying mechanism of the Thoeris defense system implies that NAD+ degradation is a unique strategy for bacterial anti-phage resistance (13).

Among these newfound systems, we focus on the Gabija bacterial defense system, which contains two components, GajA and GajB. As shown in the previous study, the Gabija system from *Bacillus cereus* VD045 showed potent defense to bacteriophage phi29, rho14, phi105 and SpBeta (10). Bioinformatic analysis suggested that Gabija is widely distributed in bacteria and archaea and exists in at least 8.5% of the sequenced genomes that are analyzed (4360 genomes) (10). As a comparison, CRISPR/Cas systems are found in about 40% of all sequenced bacteria (14,15), R-M systems are found in about 75% of prokaryote genomes (16), and prokaryotic Argonautes (pAgos) and BREX appear in about 10% of sequenced prokaryote genomes (17,18). Many known bacterial defense systems attack bacteriophage genomic DNA and most of their elements have the ability for specific nucleic acid processing, such as CRISPR/Cas and R-M systems (4,5). In this study, we aim to elucidate the function of GajA protein, which consists of an N-terminal ATPase-like domain and a C-terminal TOPRIM domain and was predicted as an ATP-dependent nuclease. Among characterized nucleases, the recently reported nonspecific nucleases of the OLD family involved in DNA repair and replication, including BpOLD, XccOLD and TsOLD (19,20), share the highest homology with GajA. GajB was predicted as a UvrD helicase and in some bacterial genomes the GajB homolog is absent thus the GajA homolog is the sole component of potential Gabija (10,19,20).

In this work, we purify and characterize the function of GajA in detail. We found that GajA exhibits specific endonuclease activity. We define the cleavage site, recognition sequences, optimal reaction conditions and functional domains of GajA. We further revealed that GajA activity is negatively regulated by nucleotides and the H320A mutation in the ATPase-like domain relieves the inhibition of ATP to GajA cleavage activity. Together we demonstrate that GajA is a novel nucleotide-sensing endonuclease and proposed a novel strategy relying on one protein for anti-phage resistance as the molecular mechanism underlying the Gabija bacterial defense system.

## MATERIALS AND METHODS

### Materials

Oligonucleotides and primers were obtained from Genscript Company. The Gibson assembly kit, Quick Blunting™ Kit, Alkaline Phosphatase and T4 DNA ligase were from New England BioLabs. T-plasmid was from Vazyme Biotech. PrimeSTAR Max DNA Polymerase was from TaKaRa. DNA purification kit was from Axygen. Ni-NTA resin was from Qiagen. Preparative Superdex S200 for gel filtration was from GE Healthcare.

### Cloning, expression and purification of GajA

The predicted coding sequence for GajA (residues 1-578), its active site mutants and GajA-CTR (residues 348-578) were cloned into pET28a vector harboring N-terminal 6×His tag with Gibson Assembly Cloning Technology (21). Constructs were transformed into *E. coli* BL21(DE3) cells, which were cultured in 2 L LB medium containing 50 μg/ml kanamycin at 37°C for three hours to an OD_600_ of 0.6-0.7, and then induced with 0.1 mM IPTG for 18 hours at 12°C.

The cells were harvested and resuspended in lysis buffer (20 mM Tris-HCl, pH 8.0, 300 mM NaCl, and 0.5 mM DTT), then lysed by ultrasonication. Supernatant was collected after centrifugation for 1 hour at 13000 rpm, 4°C and filtrated with 0.45 μm filter, loaded onto Ni-NTA agarose column pre-equilibrated with 10 volumes of elution buffer (20 mM Tris-HCl pH 7.5, 300 mM NaCl), then the column was washed with 10 volumes of elution buffer containing 20 mM and 50 mM imidazole, respectively. The majority of GajA was eluted off the column by elution buffer containing 100 mM imidazole. Collected elutes were concentrated to 2.5-3 ml by Millipore Amicon Ultra-15 (30,000 MWCO) and further purified by gel filtration chromatography on a 200 ml preparative Superdex S200 column. Fractions containing pure GajA were concentrated again. Finally, GajA was dialyzed against a storage buffer containing 50 mM Tris-HCl pH 7.5, 100 mM NaCl, 1 mM DTT, 0.1 mM EDTA, 50% glycerol and 0.1% Triton X-100. Protein concentration of GajA was determined by Bradford protein quantitative kit (Bio-rad), and protein purity and concentration were analyzed by 10% SDS-PAGE stained with Coomassie blue (Bio-rad).

Active site mutations and GajA-CTR were introduced via Gibson Assembly method (21) (primers for cloning listed in Table S2) and mutants were expressed and purified with the same procedure as detailed above.

### DNA cleavage assays

To explore the requirement of GajA for metal ions, DNA cleavage experiments were carried out at 37°C in the reaction buffer (20 mM Tris–HCl, pH 9 and 0.1 mg**/**ml BSA) supplemented with 5 mM MgCl_2_, MnCl_2_, CaCl_2_, ZnCl_2_, CoCl_2_, or NiCl_2_. After screening the optimal reaction conditions, 125 ng of DNA substrate was incubated with 0.2 µM protein in a final volume of 10 µl in DNA cleavage buffer (20 mM Tris–HCl, pH 9, 1 mM MgCl_2_ and 0.1 mg**/**ml BSA). Reactions were performed at 37°C for 2 or 5 min and then stopped by addition of 2 µl 6× loading dye containing 20 mM EDTA. Samples were analyzed via native agarose gel electrophoresis. After ethidium bromide staining, the signal of the initial DNA substrate was measured and quantified using the ImageJ software. To determine the ratio of DNA degradation, the intensity of the intact DNA substrate band in each gel lane was compared to the intensity of intact DNA band in the protein-free control lane. The quantification bar graphs represent the average of three independent trials with error bars representing the standard error of the mean.

### Determination of the DNA cleavage site

A 955-nt DNA fragment derived from the lambda phage genomic DNA (λ955) was cut into two small fragments, λ372 and λ583 by GajA. The latter two fragments were recovered, blunted with Quick Blunting™ Kit and inserted into T-plasmid, respectively. Several colonies each were selected for DNA sequencing and the GajA cleavage site was deduced from the sequencing results.

### Determination of the recognition sequence

In order to identify the full recognition sequence of GajA, we inserted the DNA sequences surrounding the GajA DNA cleavage site into pUC19 plasmid and introduced various sequence alterations into the inserted region with Gibson Assembly method. The constructed plasmids were verified by DNA sequencing and DNA fragments for GajA cleavage substrates were PCR amplified using primers pUC19-F/R (Table S1).

### Synthetic DNA substrates

The synthetic DNA substrate was prepared by mixing equimolar amounts (20 µM) of complementary 56-nt oligonucleotides in a total volume of 20 µl of annealing buffer (10 mM Tris-HCl pH 7.4, 50 mM NaCl). Complementary oligonucleotides were annealed by heating at 95°C for 5 min and then gradient cooling to room temperature over a 100 min period. In the oligoduplex cleavage assay, the reaction mixtures containing 0.4 µM GajA and 0.8 µM DNA were incubated at 37°C for 2 min. The reactions were stopped by addition of the loading dye containing 20 mM EDTA and analyzed by 12% PAGE.

### ATPase assays

ATPase activity was characterized by thin layer chromatography. The reaction was carried out in ATPase reaction buffer (20 mM Tris-OAc pH 7.9, 50 mM K-OAc, 10 mM Mg-OAc, 1 mM DTT) with 4 mM ATP, 12 µM 56-bp DNA (S1 in Figure 3B) and 3 µM or 6 µM protein at 37°C. After 2 min incubation, 1 µl samples were spotted onto a polyethyleneimine cellulose TLC plate and developed with a solution containing 1 M formic acid and 0.8 M LiCl as previously described (22).

**Figure 3.**
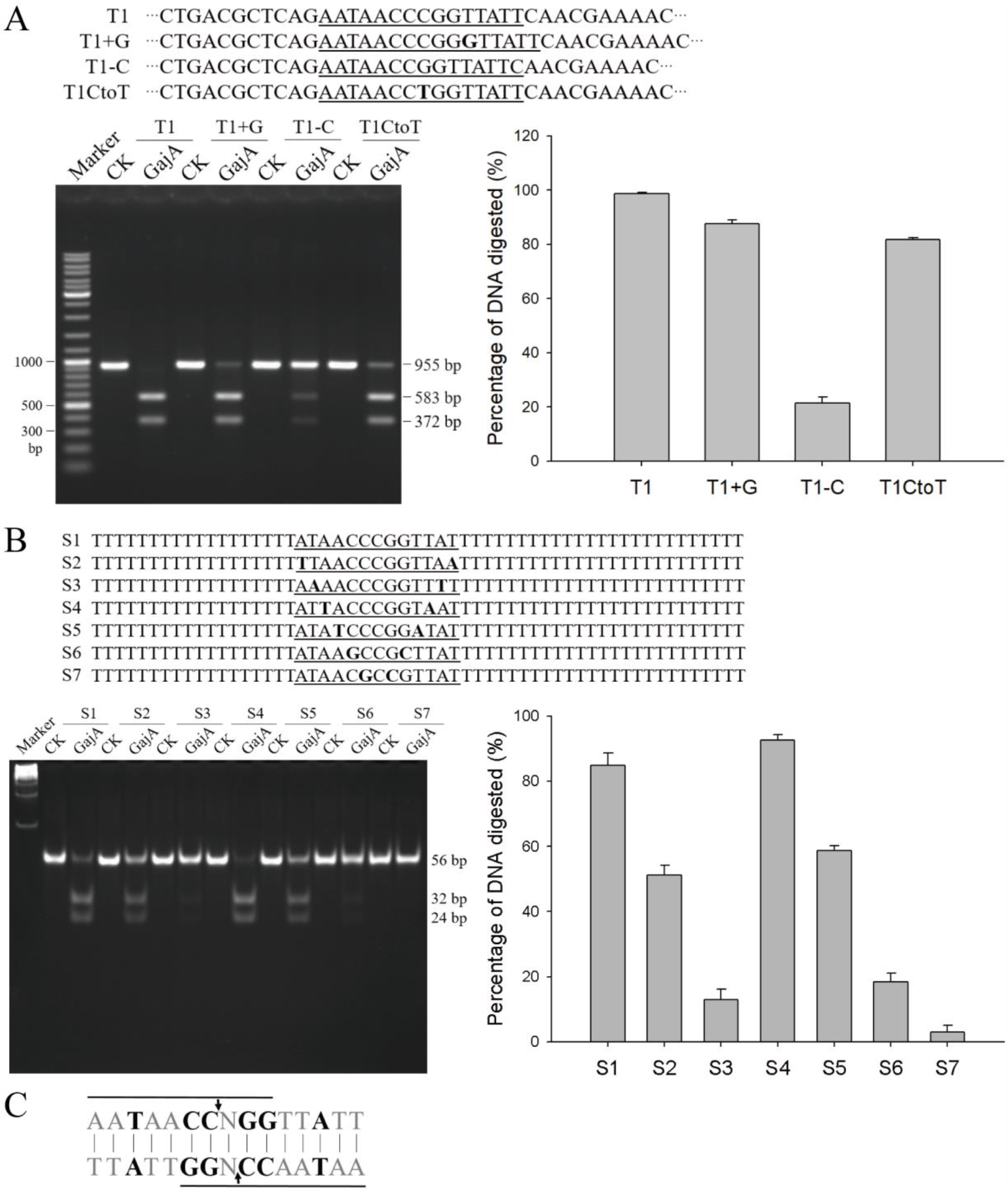
Characterization of the full optimal recognition sequence of GajA. (**A**) Cleavage efficiency (quantified as the reduction of initial DNA substrates) of GajA on variety recognition sequences. T1 is the initial λ955 DNA substrate containing the full recognition sequence (underlined) of GajA. T1+G denotes a G (bold) addition into T1 and T1-C denotes a C deletion from T1, which turn the underlined recognition sequences into complete palindromic sequences. T1CtoT represents a T (bold) mutation of the central C in T1 substrate. (**B**) Cleavage efficiency (quantified as the reduction of initial DNA substrates) of GajA on DNA oligos with base-switching in the palindromic region. The synthetic DNA substrates were prepared by mixing equimolar amounts of complementary 56-nt oligonucleotides. The DNA substrate (800 nM) was digested by GajA (400 nM) and results were analyzed by 10% PAGE. (**C**) The optimal recognition sequence of GajA. The black arrows indicate GajA cleavage sites. Lined sequences represent the minimum recognition sequence. Sequences in black are strict while those in gray are less strict for GajA recognition.

ATPase activity was quantified using PiColorLock™ phosphate detection system kit (Expedeon) that monitored the amount of free phosphate released. The reactions were performed in ATPase reaction buffer as mentioned above with 0.5 mM ATP, 2 µM 56-bp DNA (S1) and 1 µM protein for one hour at 37°C. Subsequent processing was according to the kit manual and samples were measured by NanoPhotometer® (Implen) at 650 nm.

## RESULTS

### GajA exhibits specific cleavage activity *in vitro*

The Gabija system exists in about 8.5% of all sequenced bacteria and archaea (10). In most cases it consists of two components, GajA and GajB, while in some cases the GajB predicted as a helicase is absent (10,19,20). Bioinformatics analysis indicated that GajA contains an ATPase-like domain (residues 1-341) and a TOPRIM domain (residues 348-510) (Figure 1A). As the major or sole element in the Gabija system for phage resistance, the GajA was initially suspected to function as a nuclease. To elucidate its function, GajA with over 90% homogeneity was purified as N-terminal His-tagged protein (Figure 1B). The potential nuclease activity of GajA was tested on various nucleic acid substrates and among random DNA and RNA substrates, specific cleavage activity was detected only on λDNA. A specific fragment about 2.4 kb was produced when λDNA was treated by GajA (Figure 1C). Subsequently, λDNA was divided into eight segments by PCR-amplification, among which only the 5’ foremost 6 kb region of λDNA was recognized by GajA and cleaved into a 2.4 kb and a 3.6 kb fragment (Figure S1A). This substrate was further shortened to 955 bp, which was cut into a 583 bp (λ583) and a 372 bp (λ372) fagement by GajA (Figure S1B).

**Figure 1.**
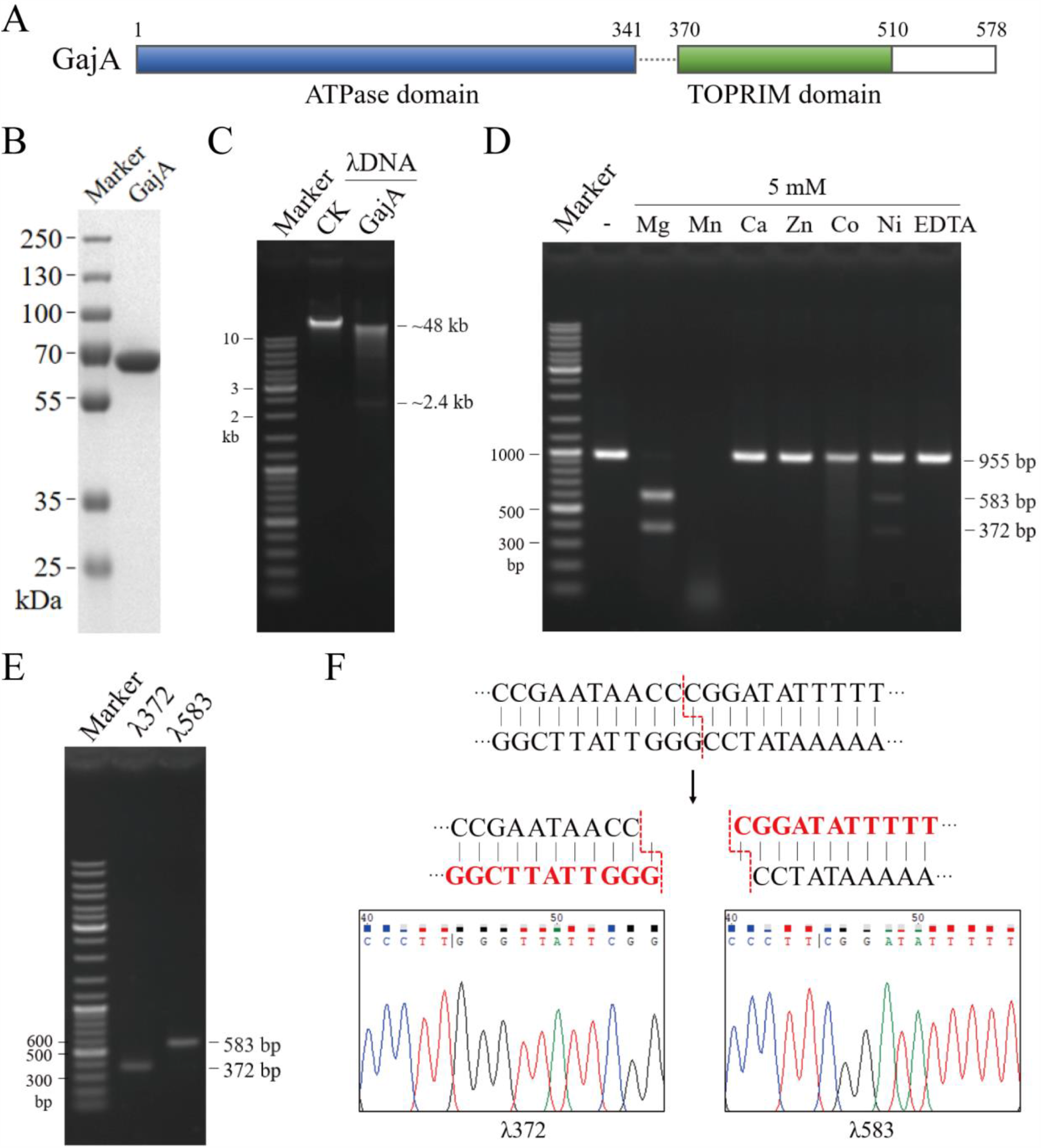
Purified GajA as an endonuclease. (**A**) Domain architecture of GajA protein. (**B**) SDS-PAGE gel showing purified GajA (69 kDa including an N-terminal His-tag). (**C**) Cleavage of linear λDNA by GajA. (**D**) GajA nuclease activity is dependent on metal ion co-factors. The reaction mixtures containing 20 mM Tris-HCl pH 9, 0.1 mg**/**ml BSA, 20 nM λ955 DNA, 200 nM GajA and 5 mM metal ion (MgCl_2_, MnCl_2_, CaCl_2_, ZnCl_2_, CoCl_2_, or NiCl_2_ as shown on top of gel) were incubated at 37°C for 5 min. Gel bands corresponding to the 955-bp λ955 DNA substrate, a 583-bp and a 372-bp DNA products resulted from specific endonuclease cleavage were annotated. Reactions with no metal ion added (-) and 5 mM EDTA were included as control. (**E**) Recovery of the two DNA fragments (λ372 and λ583) from GajA endonuclease cleavage for cloning and sequencing. (**F**) GajA cleavage site and pattern determined according to DNA sequencing results. The λ372 and λ583 fragments were inserted into T-plasmid and the regions nearby the cleavage site were sequenced, respectively. Red sequences were the terminal sequences of each fragment derived from sequencing results (bottom) and the dotted red lines demonstrate the cleavage site.

With λ955 DNA as substrate, we first examined the effect of divalent cations on the efficiency and specificity of GajA (Figure 1D). Cleavage efficiency was quantified by comparing the band intensity in each lane and calculating the percentage of DNA digested relative to the control. At the same divalent cation concentration of 5 mM, GajA exhibits rapid specific cleavage in the presence of Mg^2+^, degrading approximately 100% of the substrate within 5 minutes. In the presence of Mn^2+^, GajA degrades the DNA substrates into small pieces without showing specificity. Weak but specific GajA activity was observed in presence of Ni^2+^ while weak and nonspecific cleavage was shown in the presence of Co^2+^. Ca^2+^ and Zn^2+^ have no prominent influence on GajA activity (Figure 1D). These data indicate that metal ions are required for GajA activity and Mg^2+^ is optimal for the specific DNA cleavage of GajA. Thus, the λ583 and λ372 fragments resulted from the specific cleavage by GajA in the presence of Mg^2+^ were purified (Figure 1E) and cloned into T-plasmid, respectively, to reveal the cleavage site of GajA. 15 monoclones each were selected for DNA sequencing and the results were uniform, through which the GajA cleavage site was deduced (Figure 1F). Apparently, the cleavage site does not locate in a typical palindromic sequence as those recognized by restriction enzymes.

### The optimal reaction condition of GajA

Before investigation on the GajA recognition sequence, we optimized the reaction conditions including concentration of metal ions, pH and temperature for GajA activity. GajA cleavage activity is optimal in the presence of 1-5 mM Mg^2+^ and decreases at higher or lower concentrations of Mg^2+^ (Figure S2A). GajA exhibits specific DNA cleavage at low concentrations of Mn^2+^ (Figure S2B), with [Mn^2+^] higher than 20 µM nonspecific cleavage appears. Calcium does not potentiate and instead inhibit GajA nuclease activity when supplied with magnesium (Figure S2C). In 20 mM Tris-HCl, GajA cleavage is most efficient at pH 9 (Figure S3A) and 37°C or 42°C (Figure S3B). GajA is sensitive to salt as 100 mM NaCl or KCl completely inhibit its activity (Figure S4). Therefore, we have established the optimal reaction condition for GajA (20 mM Tris-HCl pH 9, 1 mM MgCl_2_ and 1 mM DTT, at 37°C). With this condition the GajA exhibits rapid DNA cleavage activity, as over 60% of the initial 20 nM DNA substrate was digested by 200 nM GajA after 30 seconds, and over 96% DNA substrate was digested after 120 seconds (Figure S5).

λ955 DNA fragments, whether prepared by PCR amplification or directly extracted from plasmid, were cleaved by GajA with similar efficiency (Figure S6), indicating that GajA recognition is not dependent on DNA modification. GajA also cuts the ‘supercoiled’ pUC19 plasmid containing the λ955 sequence into ‘linearized’ and ‘nicked’ DNA (Figure S6). GajA exhibits no nuclease activity on single-stranded DNA, double-stranded RNA or single-stranded RNA (Figure S7). These data suggest that GajA cuts dsDNA specifically in a sequence-dependent manner.

### Recognition sequence of GajA

The λ955 DNA fragment was cut into a 583 bp (λ583) and a 372 bp (λ372) fragment by GajA (Figure S1B). However, neither T-plasmid inserted with λ583 or λ372 is cleaved by GajA, suggesting that sequences on both sides of the cleavage site are required for GajA recognition. Therefore, we focused on the sequences surrounding the cleavage site in λ955 and gradually shortened it to the minimum GajA recognition sequence. Various recognition sequences were constructed into the pUC19 plasmid and then amplified by the common primers to be tested as GajA substrates. First we found that a 16 bp sequence 5’-GAATAACCCGGATATT-3’ containing the cleavage site is sufficient for GajA recognition (Figure S8). With gradual one-by-one nucleotide shortening on either side of the 16-bp sequence, GajA cleavage efficiency gradually decreases correspondingly (Figure 2A and B), until reaching into the center CCCGG sequence, of which a single nucleotide deletion on either side abolishes GajA activity. Meanwhile, truncations on both ends of the 16-bp sequence simultaneously caused rapid decline of GajA cleavage efficiency (Figure 2C). From these results, it seemed that the center 5-bp GC-rich core sequence in combination with the 5-bp AT-rich wing sequence on either side (AATAACCCGG or CCCGGATATT) is a minimum recognition sequence to maintain GajA cleavage, while the overlapping of these two minimum recognition sequences constitutes the full 15-bp restriction site for GajA. Interestingly, we noticed that in the 15-bp GajA restriction sequence (5’-AATAACC*C*GG**A**TATT-3’) as deduced above, the 7-bp sequences on both sides of the center cytidine (italic) are nearly palindromic. An A (bold) to T mutation to perfect the palindrome resulted in the most efficient GajA restriction sequence (5’-AATAACC*C*GG**T**TATT-3’) (Figure S9).

**Figure 2.**
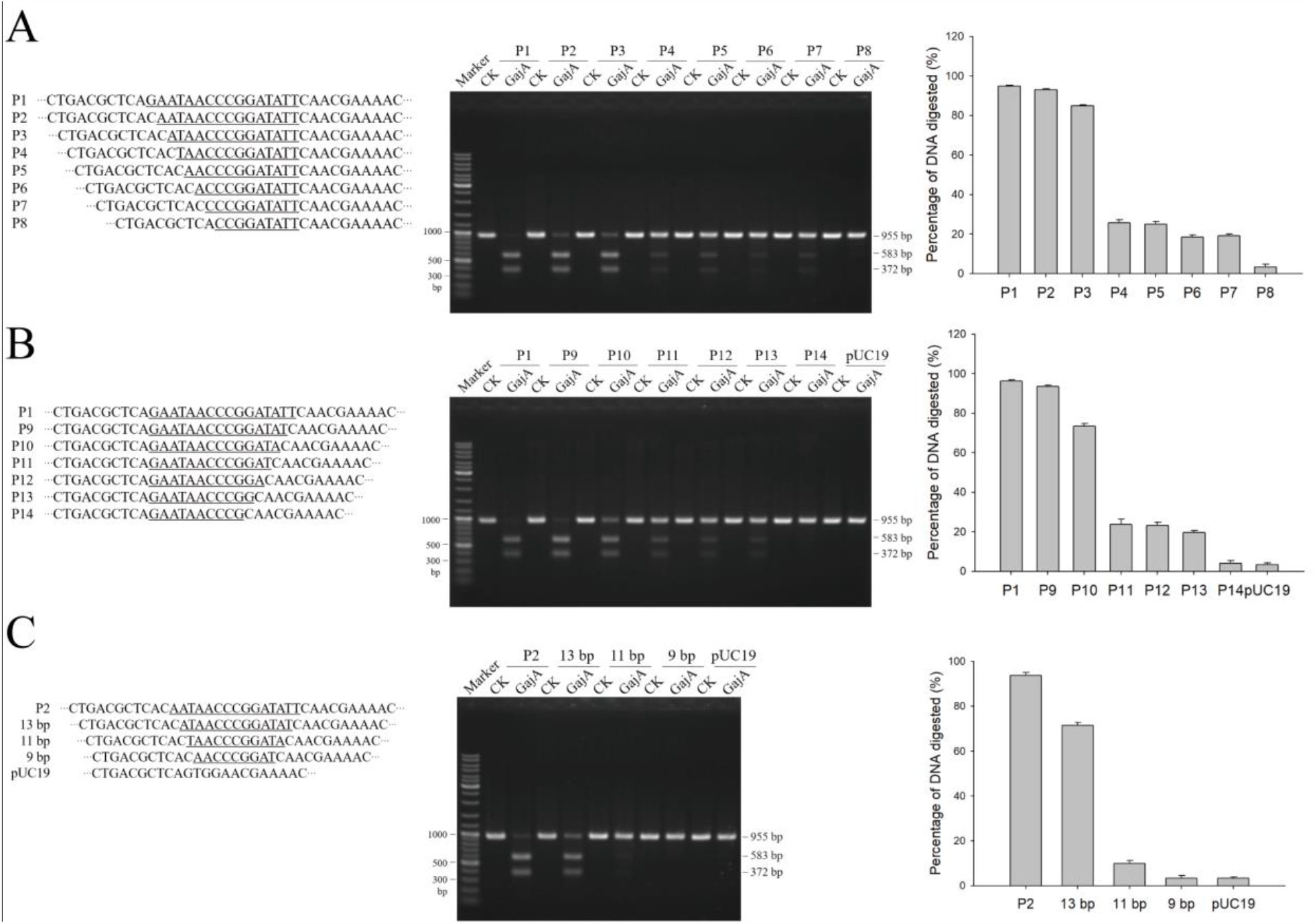
Screening for the minimum GajA recognition sequence. The preliminary recognition sequence of GajA was shortened one by one nucleotide from the left end (**A**), right end (**B**), or both ends (**C**) and DNA fragments containing the resulted sequences were PCR-amplified as substrates to test GajA cleavage efficiency. DNA digested was quantified using ImageJ software as described in Materials and Methods. All graphs represent the average of three independent trials with error bars representing the standard error of the mean.

We examined the degeneration of the GajA recognition sequence. Either a G addition or a C deletion in the core region to make the whole sequence palindromic reduced the cleavage efficiency (Figure 3A). Conversion of the center C to T only decreased the cleavage slightly (and the effect of G or A was concurrently examined at the equal position in the complementary DNA strand), thus the center nucleotide can be any of the four bases (Figure 3A). We also switched each pair of nucleotides in the palindromic region and found that all the GC-rich core sequence (except for the center nucleotide) and the middle nucleotide of the AT-rich wing sequence are more crucial for GajA recognition, as alterations of these nucleotides decreases the GajA cleavage activity more severe than those of other positions (Figure 3B). Altogether, we present the full recognition sequence and cleavage pattern for GajA as a restriction endonuclease (Figure 3C).

### GajA may function as a restriction enzyme or a site-specific nicking enzyme

The GajA restriction sequence (5’-AATAACCNGGTTATT-3’) are constituted of two overlapping (5’-AATAACCNGG-3’) sequences of opposite orientation (Figure 3C). And above results showed that disruption in one of the two overlapping sequences decreases but not abolishes the activity of GajA (Figure 2A and B), leading us to suspect that each of the 5’-AATAACCNGG-3’ sequences function independently to cause a nicking activity of GajA. As single nicking event in linear dsDNA is not easy to observe, we constructed plasmid P1 and P13. P1 contains the intact GajA restriction sequence thus the suspected nicking recognition sequence 5’-AATAACCNGG-3’ are presented in both DNA strands and orientations, while P13 contains only a single 5’-AATAACCNGG-3’ sequence (Figure 4A). After GajA treatment, the majority of supercoiled P1 plasmid was cleaved into linearized DNA, with a small portion of nicked plasmid. In contrast, most of P13 plasmid was turned into nicked DNA (Figure 4A and B). The overall cleaved portion for both plasmids is similar. These results confirmed that the GajA is a nicking endonuclease and a 5’-AATAACCNGG-3’ sequence is sufficient to trigger its DNA nicking activity. The unique organization of the recognition sequence renders GajA functional flexibility, the endonuclease may function as a restriction enzyme or a site-specific nicking enzyme, depending on the arrangement of the recognition sequence (Figure 4B bottom).

**Figure 4.**
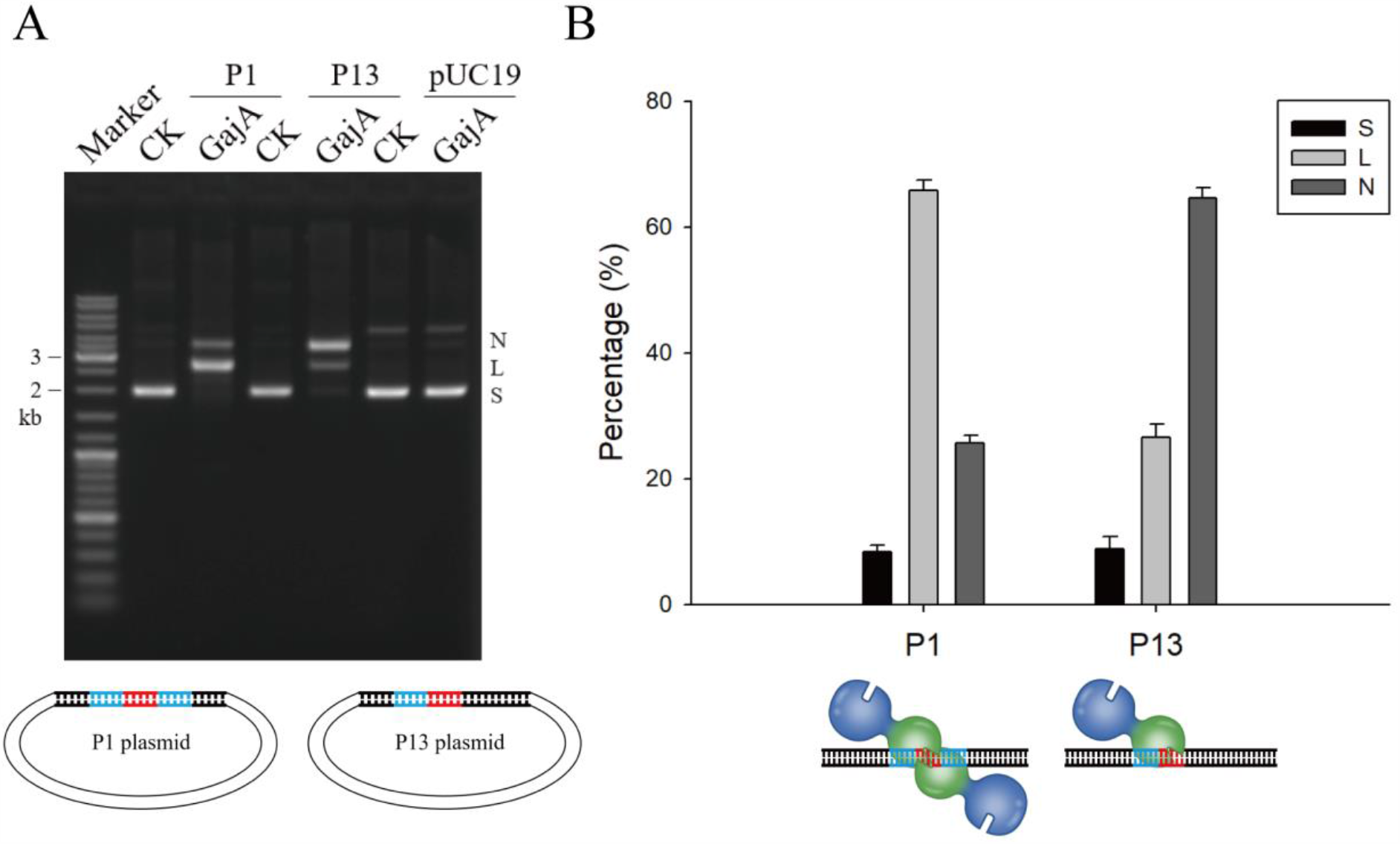
GajA may function as a restriction enzyme or a site-specific nicking enzyme depending on the organization of recognition sequences. (**A**) GajA cleavage patterns on various plasmids. P1 plasmid contains the complete GajA restriction sequence consists of two overlapping minimum recognition sequences; P13 plasmid contains one minimum recognition sequence; and pUC19 plasmid without GajA recognition sequence was used as a control. ‘N’, ‘L’, and ‘S’ denote the positions of gel bands corresponding to ‘nicked’, ‘linearized’, and ‘supercoiled’ DNA, respectively. In the plasmid diagram, the core GC-rich region of the GajA recognition sequence was shown in red and the AT-rich wing region in blue. (**B**) Quantification of products of GajA cleavage on plasmid P1 and P13. Proportion of nicked, linearized, and supercoiled DNA after GajA treated were compared. Bottom diagram depicts the two action modes of GajA endonuclease.

### The cleavage activity of GajA is suppressed by nucleotides

A N-terminal ATPase-like domain occupies more than a half of the GajA polypeptide (Figure 1A), naturally raising the possibility that it hydrolyzes ATP to stimulate the endonuclease activity as reported previously for MLH1-MLH3 complex (23). However, thin layer chromatography ATPase assay revealed that GajA has no ATPase activity, which is further confirmed by monitoring the amount of free phosphate released (Figure S10A and B). When we added ATP into GajA reaction to test whether it enhances the endonuclease activity, we surprisingly found that GajA activity was severely inhibited by ATP or the nonhydrolyzable analog AMP-PNP, 1 mM ATP or AMP-PNP fully suppresses GajA activity (Figure 5A). We further tested the effect of all sets of NTPs and dNTPs, including ADP and AMP, on GajA activity. The results show that GajA activity is strongly inhibited by all NTPs and dNTPs, even ADP, while AMP shows no significant influence (Figure 5B). Apparently, the endonuclease activity of GajA is negative-regulated by nucleotides and the regulation is not dependent on the hydrolysis of ATP or other tri-phosphates.

**Figure 5.**
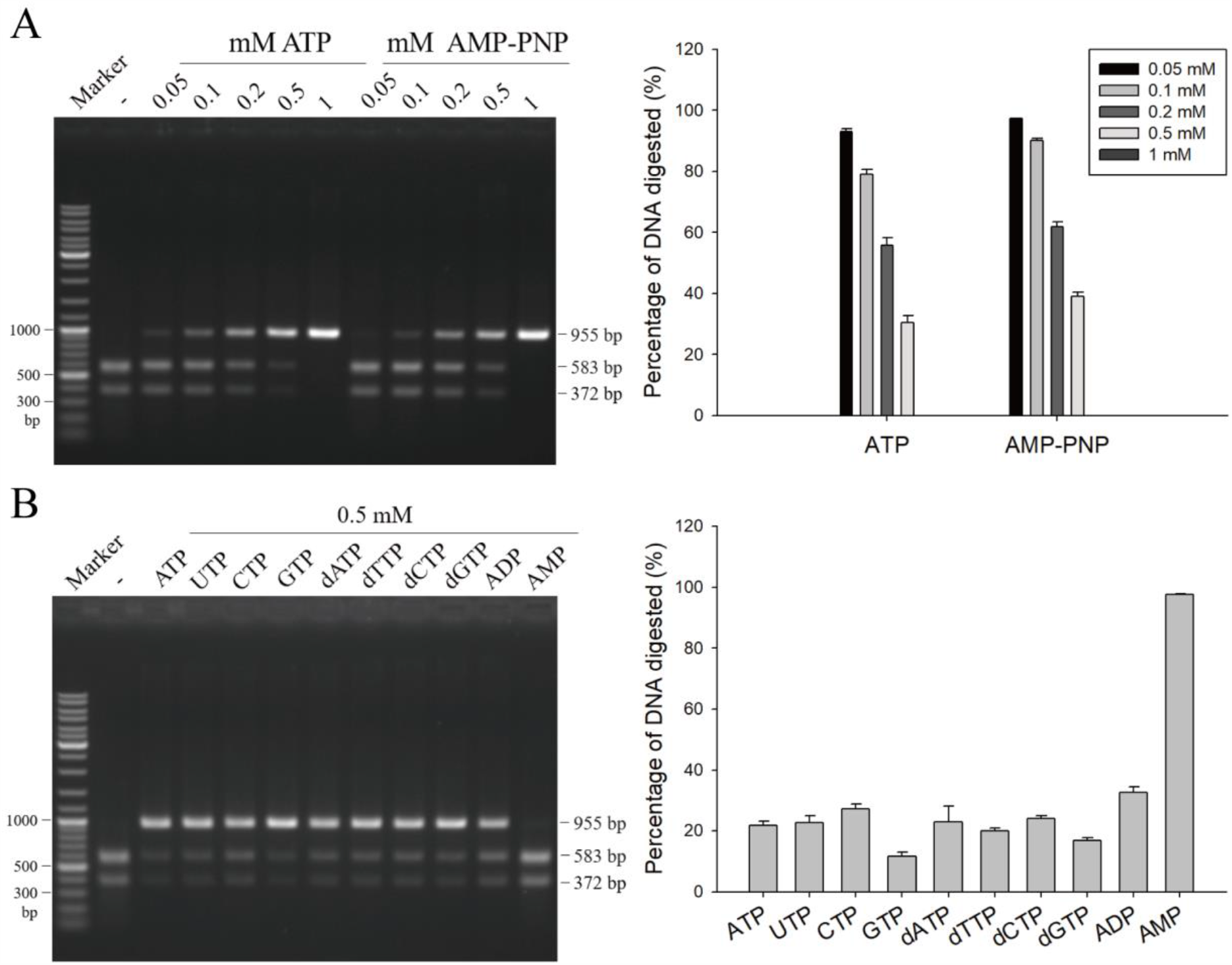
The endonuclease activity of GajA is inhibited by nucleotides. (**A**) Representative gel and quantification of GajA endonuclease activity on λ955 DNA in the presence of increasing amounts of ATP and AMP-PNP. (**B**) The effect of NTP, dNTP, ADP and AMP on GajA endonuclease activity. All reactions contain 20 nM λ955 DNA and 200 nM GajA and were incubated at 37°C for 5 min. Lanes labeled with dashes indicate no nucleotide addition. Initial DNA digested was quantified using ImageJ software. Bar graphs represent the average of three independent experiments with error bars representing the standard error of the mean.

### GajA functional domains

Based on sequence homology, the OLD family proteins are the closest to GajA among characterized nucleases (Figure S11). BpOLD and TsOLD also consist of an ATPase domain and a TOPRIM domain, resembling the domain organization of GajA (19,20). As previously reported, the ATPase domain of TsOLD is functional in ATP hydrolysis and the TOPRIM domain carries out non-specific nuclease activity (20). Despite the functional discrepancy between the OLD proteins and GajA, we seek to identify key residues of the GajA active site based on their sequence similarity. Meanwhile we performed multi sequence alignment of GajA and its homologs from various species. The alignment indicated potential key residues in their active sites, such as the conserved TOPRIM glutamate and the DxD motif (Figure S11). Therefore, we performed alanine screen of some conserved residues in both the ATPase-like domain and the TOPRIM domain of GajA, constructing K35A, H320A, E379A, D511A and K541A GajA mutants (sites labeled by asterisks in Figure S11). In addition, we also constructed the N-terminal domain-truncated version of GajA (GajA-CTR), leaving only the TOPRIM domain. GajA mutants were purified with the same procedure as that for the wild-type protein (Figure 6A). Mutations of the key residues in TOPRIM domain (E379A, D511A and K541A) completely abolish the endonuclease activity of GajA, while those in the ATPase-like domain (K35A and H320A) show no effect (Figure 6B), confirming that the endonuclease active site locates in the TOPRIM domain. However, the H320A mutation in the ATPase-like domain partially relieves the inhibition of ATP on GajA endonuclease activity (Figure 6C), suggesting that ATPase-like domain mediates the regulation of endonuclease by nucleotide-sensing. Consistently, GajA-CTR exhibited no endonuclease activity (Figure 6B), implying the indispensable role of the ATPase-like domain. Although the homologous residue in TsOLD protein is crucial for the ATP hydrolysis (20), mutation of K35 does not affect the inhibition by ATP (Figure 6C), supporting the model that the binding but not hydrolysis of ATP (and other nucleotides) by the ATPase-like domain mediates the regulation of the endonuclease activity of GajA.

**Figure 6.**
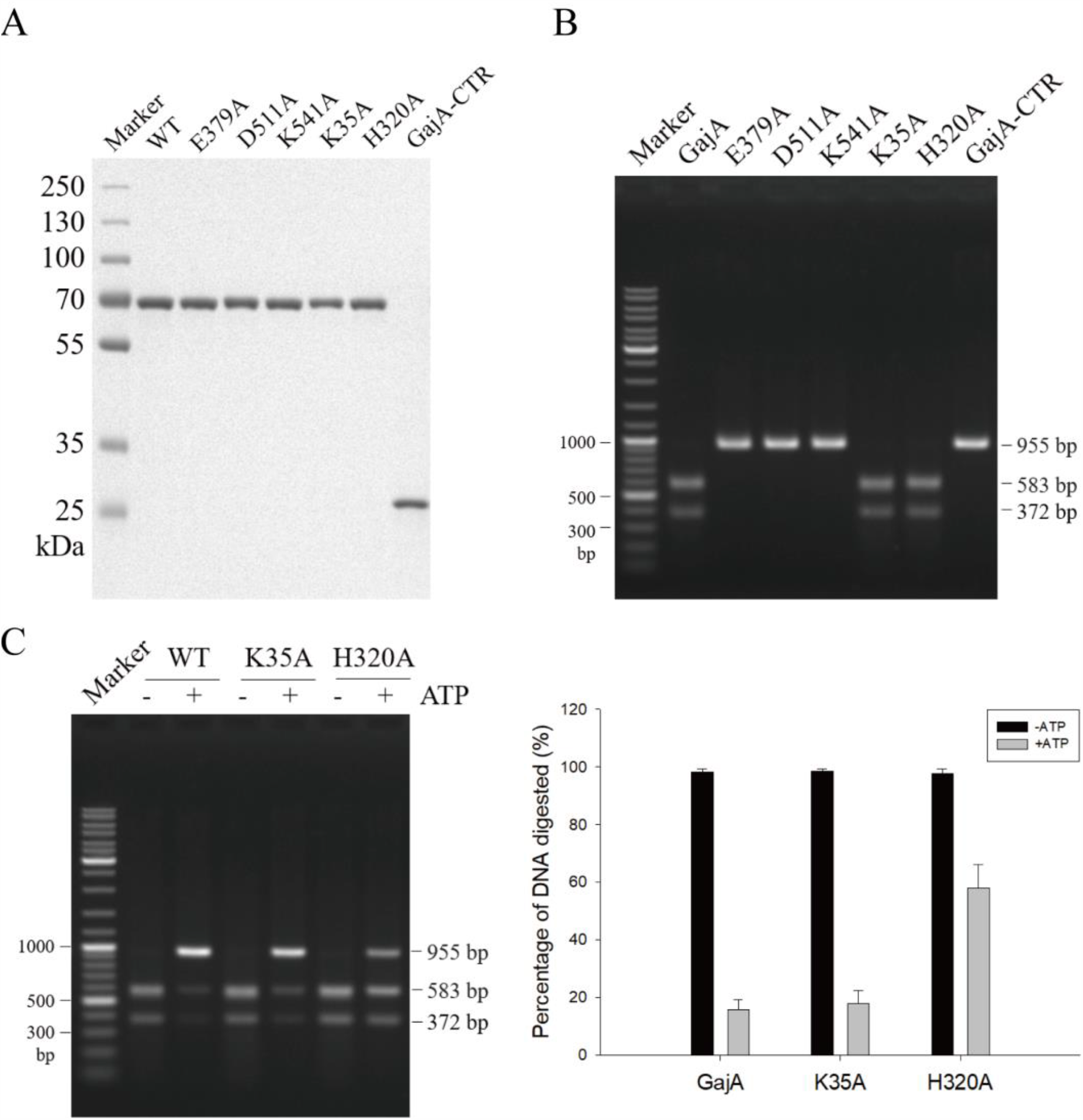
Investigation of GajA functional domains by site-specific mutagenesis. (**A**) SDS-PAGE analysis of purified wild-type (WT) GajA, GajA mutants and the C-terminal polypeptide (CTR) of GajA. (**B**) Endonuclease activity of the proteins in (A). (**C**) The effect of K35A or H320A mutation on the ATP inhibition of GajA activity. The H320A but not K35A mutation partially relieves the inhibition of ATP on GajA activity.

## DISCUSSION

### GajA is a novel endonuclease

GajA exhibits specific and metal-dependent DNA cleavage activity (Figure 1D). In contrast, as the homologs of GajA, BpOLD, XccOLD and TsOLD show non-specific nuclease acticity (19,20). Phylogenetic analysis of GajA and its homologs from different species indicates the evolutionary relationship between GajA and BpOLD, TsOLD (Figure S12). Although the OLD family proteins function in DNA repair and/or replication (19,20), while GajA is responsible for anti-phage defense, they might have evolved from common ancestor. GajA and its homologs including OLD proteins are properly conservative in the TOPRIM active sites, such as the conserved glutamate and the DxD motif, although the overall sequence homology is low (Figure S11). Like BpOLD and TsOLD with a two-metal catalysis mechanism (19,20), GajA may share the common mechanism due to the conservatism of the active sites (Figure 6B). Apparently, the sequence specificity of the OLD family proteins diminished or have been lost during functional divergency. In contrast, it is likely that the ATPase-like domain of GajA lost the ATP-hydrolysis activity as observed from OLD family proteins, instead it retained the nucleotide-binding function and has evolved into a regulatory domain. The binding of nucleotides by GajA seems not specific for the base and sugar ring, as all NTPs and dNTPs shown similar effect on GajA activity. However, the phosphate group of nucleotides play a crucial role in the binding and regulation, as AMP failed to inhibit GajA activity while ADP and ATP shown strong inhibition (Figure 5B). Further structural studies are needed to clarify the specific mechanism of GajA function.

GajA may function as a natural site-specific nicking endonuclease (NEase) upon a 10-bp recognition sequence 5’-AATAACC↓**N**GG-3’ (down arrow marks the nicking site) (Figure 4A and B). Known natural site-specific NEases were divided into two major groups: one group includes small HNH NEases from phage or prophage genomes that nick dsDNA sites with 3- to 5-bp specificities, for example, Nt.CviPII (↓CCD) originally found in chlorella virus (24,25). These small HNH NEases are involved in phage DNA packing and pathogenicity island mobility and are widespread in nature (26,27). Other phage-encoded NEases with longer recognition sequences may also be classified into this group, such as the phage group I intron-encoded HNH homing endonucleases I-PfoP3I that nicks DNA sites of 14-16 bp (28) and T7-like phage ΦI encoded I-TslI that nicks DNA sites with a 9-bp core sequence (29). Another group of NEases with 3- to 7-bp specificities are natural components of restriction systems, such as Nb.BtsI, the large subunit (B subunit) of BtsI (30). Distinct from these natural nicking enzymes, the GajA is a free-standing nicking enzyme from bacteria, relying on a TOPRIM domain to nick the DNA within the specific 10-bp sequence. And proper expansion of 10-bp nicking sequence may turn GajA into a Type II restriction endonuclease.

Type II restriction endonucleases cleave within or at short specific distances from a recognition site (7,8). They mostly require magnesium and are single function enzymes independent of methylase. At present, all characterized Type II restriction endonucleases were classified into 11 subtypes by their function modes, namely: A (recognize **<u>a</u>**symmetric sequences and cleave within, or a defined distance away from the sequence), B (cleave DNA on **<u>b</u>**oth sides of their recognition sequence, releasing a small fragment that contains the recognition sequence), C (**<u>c</u>**ombined enzymes containing endonuclease and methyltransferase activities in the same protein), E (Type IIP enzymes with allosteric **<u>e</u>**ffector domains that stimulate catalysis when bound to additional recognition sequences), F (bind two recognition sequences and cleave coordinately, hydrolyzing all four DNA strands at once), G (with a DNA-cleavage domain and a **<u>g</u>**amma-class DNA-methylation domain in a single polypeptide chain), H (**<u>h</u>**ybrid, part Type I and part Type II), M (require **<u>m</u>**ethylated recognition sequences), P (recognize **<u>p</u>**alindromic (symmetric) DNA sequences and cleave symmetrically within the sequence), S (cleavage is **<u>s</u>**hifted to one side of the sequence, within one or two turns of the double helix away) and T (act as he**<u>t</u>**erodimers, and comprise **<u>t</u>**wo different subunits) (7,31-33). Given the palindromic recognition sequence with an additional central base (bold) 5’-AATAACC**N**GGTTATT-3’, GajA acts similarly as a typical Type IIP restriction enzyme to cleave both strands within the recognition site and leave one-nt sticky ends on products (Figure 3C). However, unlike any known Type IIP restriction enzyme, when part of the full restriction sequence 5’-AATAACCCGG-3’ was supplied, GajA functions as a native nicking endonuclease to cleave one strand of DNA (Figure 4). In addition, to our knowledge GajA, as a Type II restriction enzyme is unique in that its activity is regulated allosterically by an associated nucleotide-binding domain and nucleotide-sensing. Thus, we suggest that GajA is classified as the representative of a new subtype of Type II restriction enzymes named Type IIR (**<u>r</u>**egulated).

### Proposed model for the Gabija anti-phage defense mechanism

In the preliminary genetic study, the fragment located in 93871-97763 of the *Bacillus cereus* VD045 genome, which is predicted containing GajA and GajB, is cloned into the plasmid pSG1-rfp, and then transformed into the donor bacterial strain, *Bacillus subtilis* str. BEST7003, which show strong defensive effect to various phages (10). The predicted GajA and GajB genes are separated by just one nucleotide, thus we speculated that such a cassette may only express the GajA protein but not GajB. Our overexpression of the GajA/B gene fragment with the same organization in *E. coli* confirmed that only GajA was expressed. Considering that in some cases the GajB homolog is absent (10,19,20), it is likely that the GajA is the key and even the sole component for defense function.

The physiological concentration of ATP is over 3 mM and the total nucleotide concentration is above 8.7 mM in *E*.*coli* of mid-log phase (34) and GajA activity is fully inhibited by 1 mM ATP *in vitro*. Therefore, the robust DNA cleavage activity of GajA should be strictly suppressed by NTPs and dNTPs at physiological concentrations. One can speculate that only drastic changes in cellular nucleotides concentration can activate the GajA endonuclease activity. Phages usually apply their own nucleases to degrade the host nucleic acids to supply the building blocks for their own, in this scenario the cellular concentrations of the degradation products NMPs and dNMPs might be temporarily high. Interestingly, GajA can avoid the inhibition from such nucleoside monophosphates (Figure 5B) to perform defense timely.

Altogether, our data suggested a model for the anti-phage mechanism of Gabija system (Figure 7A and B). GajA endonuclease activity is fully inhibited by nucleotides at physiological state (Figure 7A). The robust transcription and DNA replication of invaded phages deplete the cellular NTPs and dNTPs to a certain low level, which releases the allosteric suppression and activates the GajA endonuclease activity, and in turn results in cleavages of phage DNA (Figure 7B). Consistently, GajA recognition sequences were identified in related phages (Table S3). Meanwhile GajA recognition sequences also appear in the host bacterial genomes, indicating that GajA may also destruct genomic DNA of bacteria for abortive infection. The Gabija system based on a single nucleotide-sensing endonuclease represents a concise strategy for anti-phage defense.

**Figure 7.**
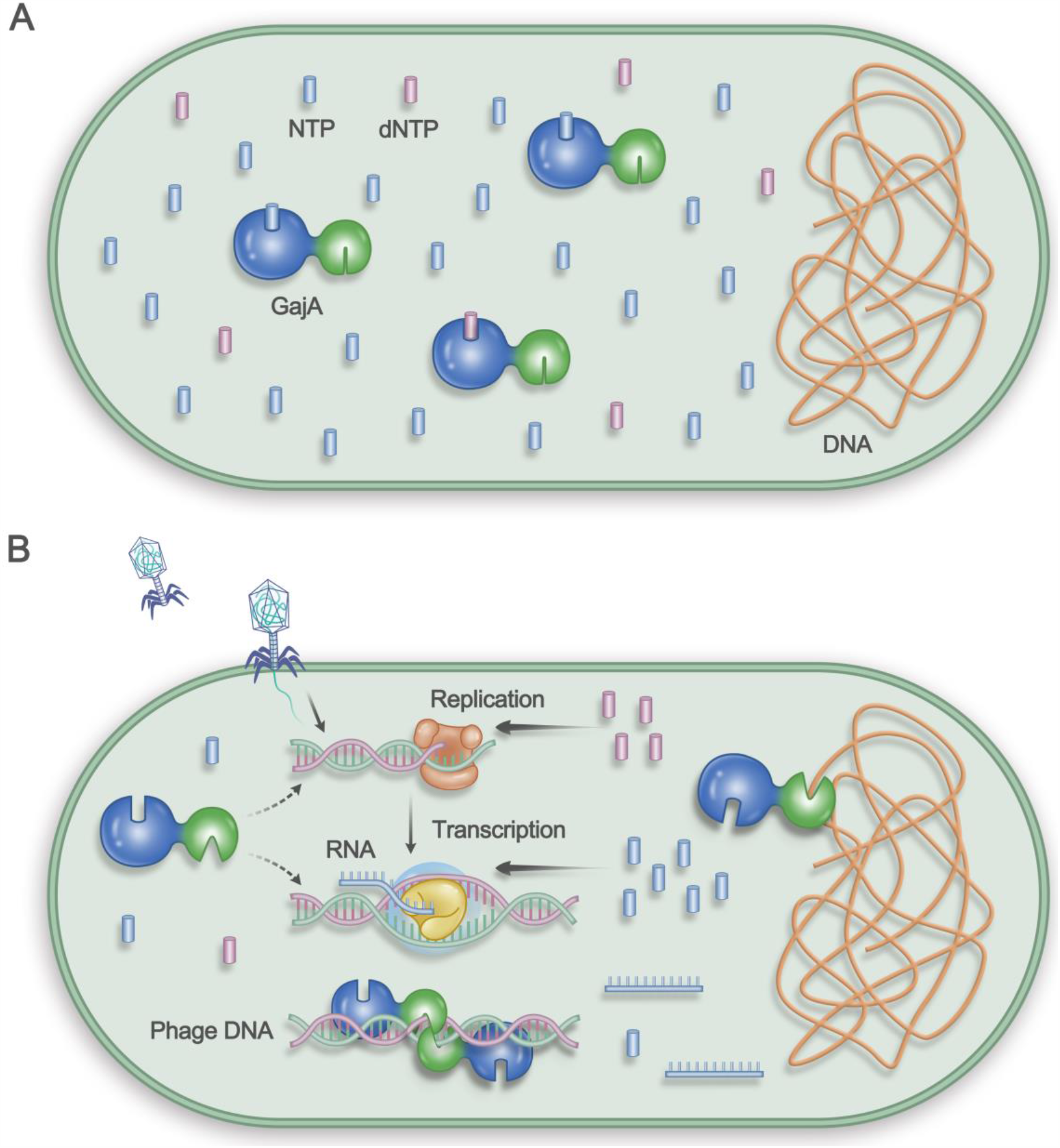
Schematic showing the proposed mechanism of Gabija anti-phage bacterial defense. (**A**) Under normal conditions, GajA endonuclease activity is fully inhibited by nucleotides at physiological concentration in bacteria. The ATPase-like domain of GajA senses and binds NTPs and dNTPs to allosterically regulates the TOPRIM domain. (**B**) During phage invasion, active phage transcription and DNA replication deplete cellular NTPs and dNTPs. When NTP and dNTP concentrations decrease to certain degree, the loss of nucleotide-binding of the GajA ATPase-like domain activates the TOPRIM domain, the latter in turn carries out phage DNA cleavage and might also mediate destruction of the own genomic DNA of bacteria for abortive infection.

## SUPPLEMENTARY DATA

Supplementary data are available online.

## ACKNOWLEDGEMENT

We thank all lab members for helpful discussion.

## AUTHOR CONTRIBUTION STATEMENT

R.C. and B.Z. conceived the project and designed the experiments. R.C. carried out the experiments. R.C., F.T.H., H.W., X.L.L., Y.Y., B.B.Y., X.L.W. and B.Z. analyzed the data. R.C. and B.Z. wrote the manuscript. All authors discussed the results and contributed to the final manuscript.

